# Snake River alfalfa virus, a tentative endornavirus infecting alfalfa (*Medicago sativa* L.) in Washington State, USA

**DOI:** 10.1101/2022.11.04.515226

**Authors:** Olga A. Postnikova, Brian M. Irish, Jonathan Eisenback, Lev G. Nemchinov

## Abstract

Here we report, for the first time, an occurrence of Snake River alfalfa virus (SRAV) in Washington state, USA. SRAV was recently identified in alfalfa (*Medicago sativa* L.) plants and western flower thrips in south-central Idaho and proposed to be a first flavi-like virus identified in a plant host. We argue that the SRAV, based on its abundance in alfalfa plants, genome structure, presence in alfalfa seeds, and seed-mediated transmission is a member of the *Endornaviridae* family.

## Main text

Snake River alfalfa virus (SRAV) was recently identified from alfalfa plants and thrips *Frankliniella occidentalis* collected in the Minidoka and Twin Falls counties of Idaho, USA (Dahan et al. 2022). Based on the genome structure and phylogeny of its RNA-dependent RNA polymerase (RdRp), the authors hypothesized that SRAV is the first flavi-like virus identified in a plant host (Dahan et al. 2022). The SRAV polyprotein, however, contained no predicted helicase domain found in all flaviviruses.

To date, no occurrences of SRAV have been reported in alfalfa or on different hosts from other locations. In this work, applying high-throughput sequencing (HTS), we detected SRAV in 50 individual alfalfa plant samples collected from 10 commercial fields in Grant County, WA. Plants used for RNA extraction displayed a diverse symptomatology that occasionally correlated with the symptoms allegedly reported for SRAV, such as yellowing and vein clearing (Fig.1), (Dahan et al. 2022). These symptoms, however, were likely due to the presence of multiple co-infecting pathogens in the same plants.

**Figure 1.**
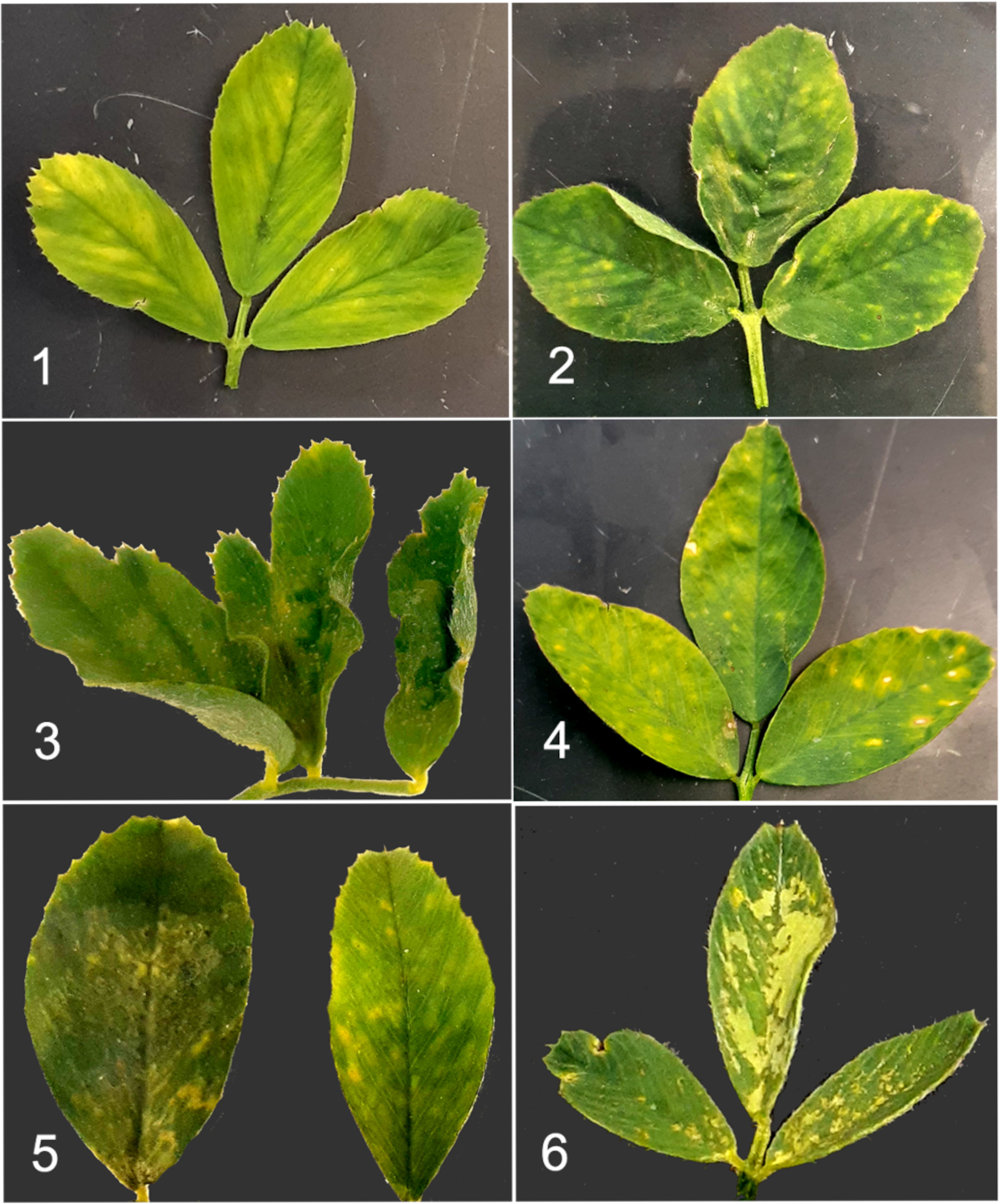
Examples of different symptomatology observed on the leaves of alfalfa (*Medicago sativa* L) plants infected with Snake River alfalfa virus (SRAV-WA1) in Washington State, USA. Sample 1 contained 7,016 reads of SRAV and 11,797 covered bases; sample 2 contained 10,684 reads mapped to the virus and 11,699 covered bases; sample 3 contained 47,175 reads mapped to the virus and 11,838 covered bases; sample 4 contained 71,267 viral reads and 11,811 covered bases; sample 5 contained 8,079 reads and 11,872 covered bases (100% vs. reference genome); and sample 6 contained 19539 viral reads and covered 11795 bases. Multiple viral, fungal and bacterial pathogens were also identified in samples in which SRAV was found. These included alfalfa mosaic virus, bean leaf roll virus, pea streak virus, partitiviruses, amalgavirus, lucerne transient streak virus, *Alternaria alternata, Alternaria arborescens, Bipolaris* spp., *Stemphylium lycopersici, Fusarium* spp., *Pseudomonas* spp., *Erwinia* spp., etc. thus emphasizing the importance of pathobiome in signs and symptoms of disease.

Total RNA was extracted using Promega Maxwell^®^ RSC Plant RNA Kit (Promega Corp., Fitchburg, WI) Library preparation was performed with Illumina TruSeq Stranded Total RNA with Ribo-Zero kit (Illumina Inc., San Diego, CA), and the sequencing platform used was HiseqX10 (PE150) (Omega Biosciences, Norcross, GA). Bioinformatic pipeline included adapter trimming followed by de-novo assembly of reads, unmapped to *M. sativa* genome and *M. truncatula* mitochondrion genomes using SPAdes (Meleshko et al. 2019) in HMM-guided mode. Phylogenetic analysis was performed with MEGA software (Kumar et al. 2016) using Maximum Likelihood method and bootstrap analysis of 1000 replicates. Conserved RdRp domains for multiple alignment were obtained using InterPro tool (https://www.ebi.ac.uk/interpro/).

All 50 unique alfalfa plant samples contained viral reads, varying in quantity from 46 to 71,267 (Table 1). Total number of reads mapping to the SRAV genome was 1,017,715 with an approximate length of each read 150 nt (Table 1). Several contigs apparently representing the complete genome of the virus were assembled de novo. The sequences varied slightly in length and contained small numbers of SNPs, indicating a presence of different genetic variants of the virus. Nevertheless, the sequence identity when compared to the reference genome (ON669064) in all cases remained >99%.

**Table 1.** Occurrence of SRAV-related reads in 10 commercial alfalfa fields (five samples per field) of Grant County, Washington State.

| Sample ID | Length | Covered % | Covered bases | Total viral reads | Reads/sample | Yield (Mb) | % >= Q30 |
| --- | --- | --- | --- | --- | --- | --- | --- |
| A2-1 | 11872 | 100 | 11872 | 8079 | 31,990,204 | 4,831 | 94.56 |
| A1-4 | 11872 | 99.9832 | 11870 | 27925 | 37,707,294 | 5,694 | 94.54 |
| A4-3 | 11872 | 99.9663 | 11868 | 22147 | 39,187,338 | 5,917 | 94.75 |
| B8-4 | 11872 | 99.9579 | 11867 | 15093 | 45,330,070 | 6,845 | 95.26 |
| A3-4 | 11872 | 99.9326 | 11864 | 44605 | 44,468,798 | 6,715 | 94.77 |
| A3-5 | 11872 | 99.7473 | 11842 | 32167 | 34,259,318 | 5,173 | 94.76 |
| B8-2 | 11872 | 99.7136 | 11838 | 47175 | 44,581,834 | 6,732 | 95.21 |
| A5-5 | 11872 | 99.6715 | 11833 | 21219 | 45,938,534 | 6,937 | 94.76 |
| B6-5 | 11872 | 99.6631 | 11832 | 13815 | 44,882,040 | 6,777 | 95.29 |
| A5-4 | 11872 | 99.6462 | 11830 | 8513 | 34,368,240 | 5,190 | 94.66 |
| B8-5 | 11872 | 99.621 | 11827 | 53101 | 37,624,098 | 5,681 | 95 |
| A1-2 | 11872 | 99.5788 | 11822 | 28724 | 42,968,950 | 6,488 | 94.74 |
| A3-1 | 11872 | 99.5115 | 11814 | 33926 | 42,255,734 | 6,381 | 94.78 |
| A2-2 | 11872 | 99.4862 | 11811 | 71267 | 37,337,248 | 5,638 | 94.71 |
| B6-1 | 11872 | 99.4862 | 11811 | 17265 | 47,716,504 | 7,205 | 93.95 |
| B7-5 | 11872 | 99.4525 | 11807 | 25870 | 31,743,182 | 4,793 | 95.13 |
| A1-5 | 11872 | 99.4441 | 11806 | 20408 | 46,898,746 | 7,082 | 94.9 |
| B6-4 | 11872 | 99.4104 | 11802 | 24605 | 36,660,924 | 5,536 | 95.13 |
| B7-4 | 11872 | 99.402 | 11801 | 29217 | 39,153,470 | 5,912 | 95.18 |
| B8-1 | 11872 | 99.402 | 11801 | 44383 | 33,674,988 | 5,085 | 94.86 |
| A1-3 | 11872 | 99.3851 | 11799 | 14704 | 34,965,248 | 5,280 | 94.65 |
| A2-5 | 11872 | 99.3851 | 11799 | 27777 | 50,933,376 | 7,691 | 94.69 |
| A4-4 | 11872 | 99.3851 | 11799 | 20253 | 31,710,774 | 4,788 | 94.62 |
| A5-2 | 11872 | 99.3851 | 11799 | 14048 | 42,018,256 | 6,345 | 94.7 |
| B10-4 | 11872 | 99.3851 | 11799 | 16305 | 35,022,936 | 5,288 | 95.1 |
| B8-3 | 11872 | 99.3851 | 11799 | 37761 | 36,163,008 | 5,461 | 95.34 |
| B9-3 | 11872 | 99.3851 | 11799 | 26815 | 41,588,106 | 6,280 | 95.12 |
| B6-3 | 11872 | 99.3767 | 11798 | 30672 | 43,616,596 | 6,586 | 95.3 |
| B9-4 | 11872 | 99.3767 | 11798 | 10475 | 50,653,058 | 7,649 | 95.3 |
| A2-3 | 11872 | 99.3683 | 11797 | 64184 | 39,012,034 | 5,891 | 94.7 |
| A4-1 | 11872 | 99.3683 | 11797 | 12057 | 35,547,054 | 5,368 | 94.6 |
| A5-3 | 11872 | 99.3683 | 11797 | 7016 | 50,538,150 | 7,631 | 94.98 |
| B6-2 | 11872 | 99.3683 | 11797 | 8332 | 55,187,384 | 8,333 | 95.25 |
| B7-1 | 11872 | 99.3683 | 11797 | 39608 | 35,432,658 | 5,350 | 95.2 |
| B7-3 | 11872 | 99.3683 | 11797 | 28372 | 45,878,018 | 6,928 | 95.27 |
| B9-2 | 11872 | 99.3683 | 11797 | 12085 | 43,347,806 | 6,546 | 95.11 |
| A2-4 | 11872 | 99.3514 | 11795 | 19539 | 32,282,770 | 4,875 | 94.76 |
| A4-5 | 11872 | 99.343 | 11794 | 4425 | 32,902,614 | 4,968 | 94.63 |
| B7-2 | 11872 | 99.343 | 11794 | 10734 | 35,619,196 | 5,378 | 95.05 |
| B10-5 | 11872 | 98.8376 | 11734 | 9244 | 35,500,288 | 5,361 | 95.15 |
| B9-5 | 11872 | 98.5428 | 11699 | 10684 | 37,603,012 | 5,678 | 95.42 |
| A3-2 | 11872 | 98.0374 | 11639 | 1288 | 39,786,654 | 6,008 | 94.56 |
| A3-3 | 11872 | 96.6139 | 11470 | 1035 | 40,166,166 | 6,065 | 94.71 |
| A1-1 | 11872 | 85.3858 | 10137 | 392 | 49,277,406 | 7,441 | 94.78 |
| B10-3 | 11872 | 41.9643 | 4982 | 83 | 41,582,718 | 6,279 | 95.19 |
| A5-1 | 11872 | 40.0775 | 4758 | 94 | 40,943,890 | 6,183 | 94.85 |
| B10-2 | 11872 | 38.0475 | 4517 | 73 | 33,886,868 | 5,117 | 94.98 |
| B9-1 | 11872 | 30.4835 | 3619 | 59 | 35,593,490 | 5,375 | 95.07 |
| B10-1 | 11872 | 26.9794 | 3203 | 46 | 37,715,730 | 5,695 | 95.36 |
| A4-2 | 11872 | 22.7257 | 2698 | 51 | 36,605,156 | 5,527 | 94.69 |

One of the de novo-assembled contigs recovered from the individual library and sample containing 8079 viral reads, was 11,811 nt in length and had 100% base coverage with the reference, thus representing a complete genome of the virus. It was 7 nt longer at the 5’ end than ON669064, which was confirmed by 5’ RACE using SMARTer^®^ RACE 5’/3’ Kit (Takara Bio USA, Inc., San Jose, CA). The 3’ end of the sequence was 59 nt longer than that of the reference ON669064 and matched another isolate of the virus, SRAV_ALF1071, found in GenBank under accession number ON669090.1 (Dahan et al. 2022). At the nucleotide level, the SRAV-WA1 (for Washington State) was 99.8% identical to the reference genome ON669064 with 18 SNPs between the two, therefore depicting an isolate of the same virus.

The genome of SRAV-WA encoded a single 3,835 amino acid (aa) polyprotein 99.9% identical to the reference and lacking a helicase domain. The results obtained by HTS were validated by RT-PCR with two sets of primers in three technical replicates. One set was ANPV_3 derived from Dahan et al. (2022), and another set of primers was designed in this work: LN1036-F, GGGAGAACCAGGAAACTGTTAG and LN1037-R, CTGTCGCATAGTCCGCTTATT. RT-PCR using both primers pairs produced correct amplicons, while no products were generated from control samples in which no SRAV-related HTS reads were found (Fig. 2). The amplicons were sequenced and validated to be SRAV.

**Figure 2.**
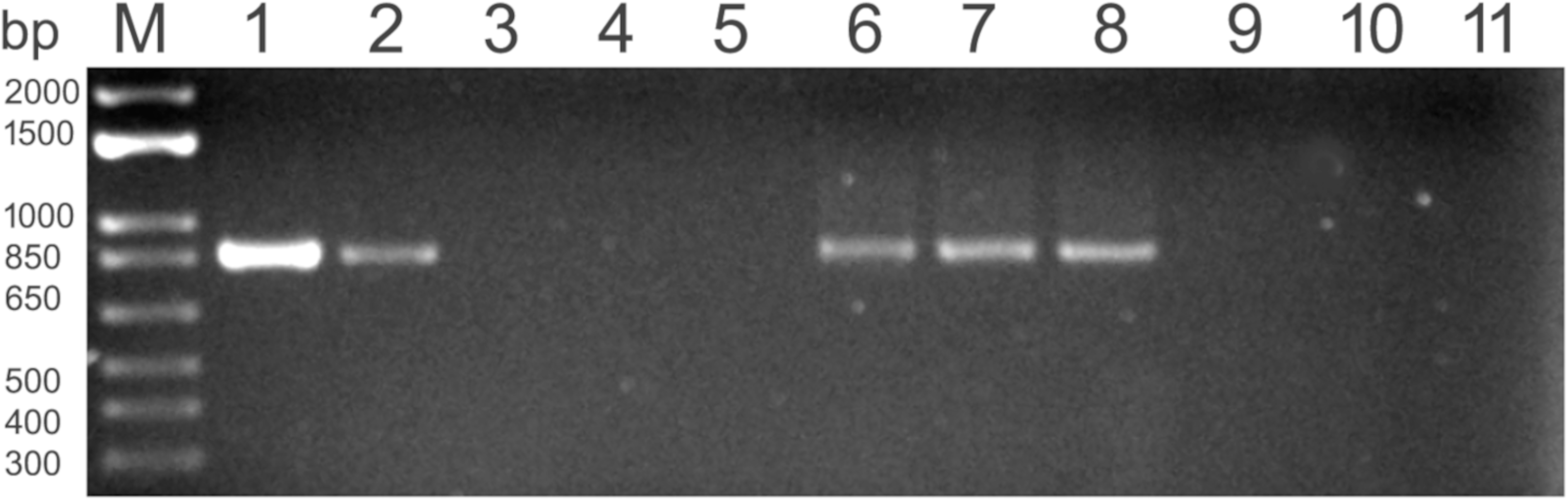
RT-PCR with primers specific for Snake River alfalfa virus. M, 1kb plus DNA marker (Thermo Fisher Scientific, Waltham, MA). Lanes 1,2: RT-PCR products amplified with primers LN1036/37 and ANPV_3, respectively. Lanes 3,4 and 5: amplification from alfalfa samples containing no SRAV reads, water control, and Taq DNA polymerase (no RT mix added to verify the absence of genomic DNA), respectively; primers LN1036/37. Lanes 6, 7, and 8: representative RT-PCR products amplified from seeds of alfalfa cultivars SW-9215, CUF101, and Maverick using LN1036/37 primers. Lanes 9,10, and 11: the same reaction controls as shown in lanes 3-5.

The ubiquitous presence of the virus in all analyzed samples indicated the possibility of persistent infection like those caused by partitiviruses and endornaviruses (Roossinck et al. 2011). Considering resemblance in the size and structure of the genome, we compared SRAV to endornaviruses. Viruses in this family infect plants, fungi, and Oomycetes, and have a linear genome of 10 to 17 kbp in length, that encodes a polyprotein ranging from 3,217 to 5,825 aa (Fukuhara, 2019; Roossinck et al. 2011; Valverde et al., 2019). Notably, several known endornaviruses, same as SRAV, lack helicase domain (Roossinck et al., 2011; Valverde et al. 2019). While members of the *Endornaviridae* family were often reported as doublestranded RNA viruses (Fukuhara, 2019; Roossinck et al. 2011), current ICTV classification describes them as single-stranded, positive-sense RNA genomes that have been characterized using replicative dsRNAs forms (Valverde et al. 2019).

Phylogenetic analysis using the polyproteins of SRAV, different viruses of the family *Endornaviridae*, and members of the family *Flaviviridae*, placed both SRAV isolates within *Endornaviridae*, in the same cluster with *Phaseolus vulgaris endornavirus 3* (Fig.3). When we performed phylogenetic analysis using InterPro-extracted RdRp domains of the *Endornaviridae* and *Flaviviriadae* (3,204-3,462 aa in SRAV-WA1), SRAV clustered with the former as well (Fig.S1). It is worth noting, however, that SRAV placement was not consistent, pointing to an irreproducibility in maximum likelihood inference and potentially incorrect phylogenies (Shen et al. 2020).

**Figure 3.**
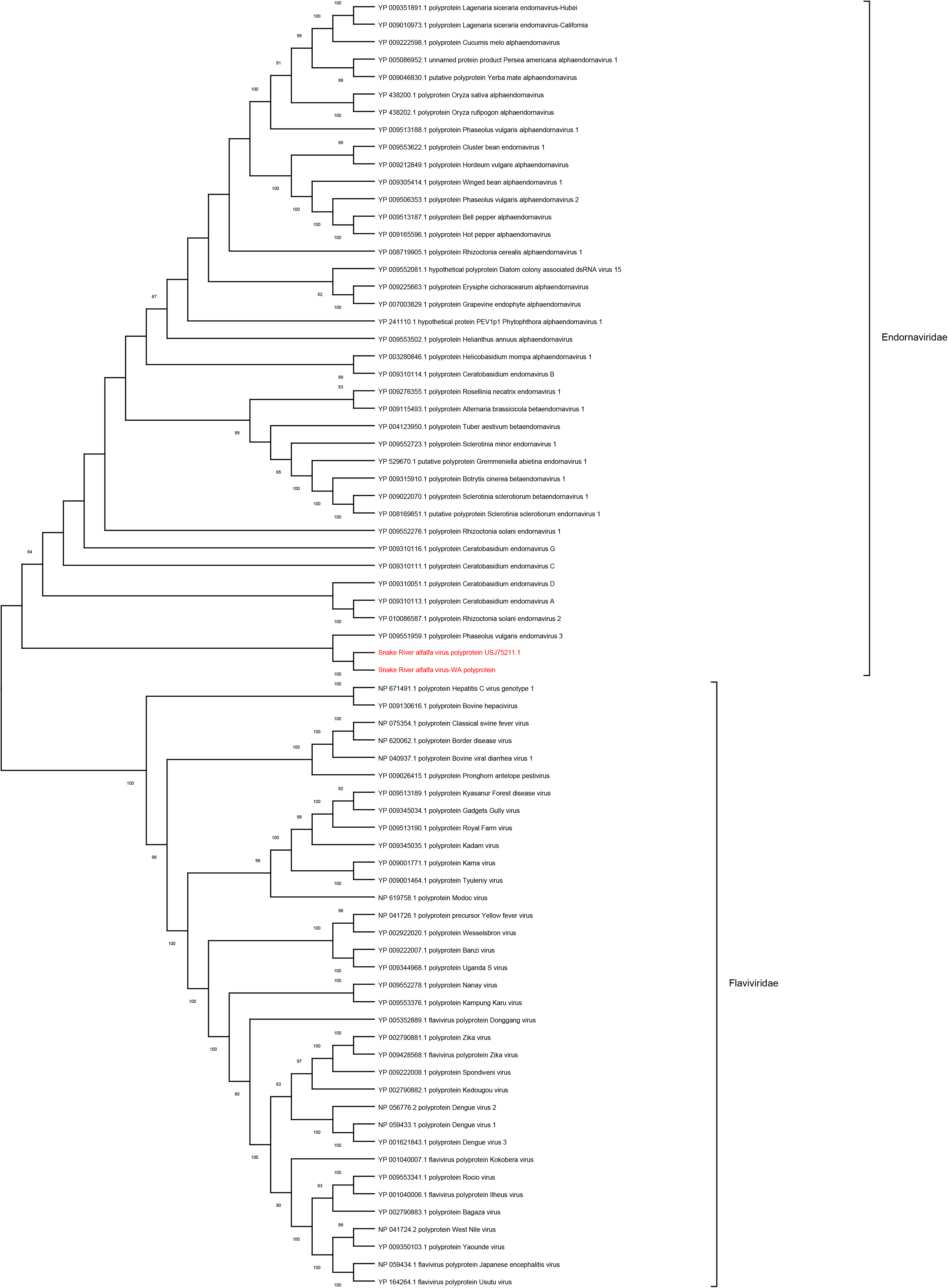
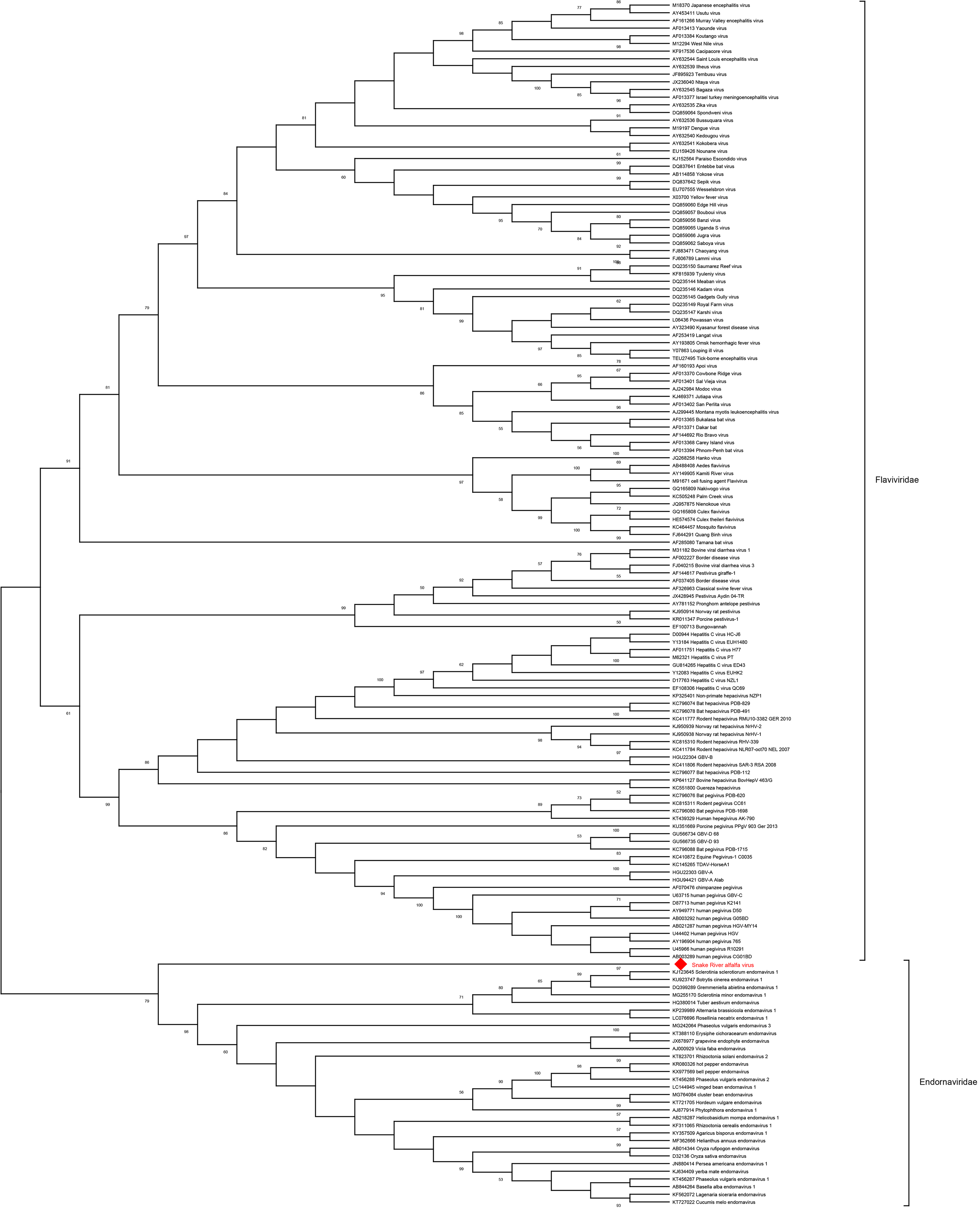
Phylogenetic relationship of SRAV with members of the families *Endornaviridae* and *Flaviviridae*. **A**. The unrooted tree was deduced from CLUSTALW alignment of the viral polyproteins and built using MEGA 11 software with Maximum Likelihood method and bootstrap analysis of 1000 replicates. **B.** The unrooted tree was deduced from MUSCLE alignment of the viral RdRP domains and built using MEGA 11 software with Maximum Likelihood method and bootstrap analysis of 1000 replicates.B

Since plant endornaviruses are transmitted through seeds via the gametes (Valverde et al. 2019), we decided to test seeds of several alfalfa cultivars for the presence of the virus by RT-PCR. Seeds were scarified with concentrated H_2_SO_4_, surface-sterilized with 70% ethanol, and rinsed with sterile water (Nemchinov and Grinstead, 2021). Total RNA was extracted with Takara Plant and Fungal RNA isolation kit (Takara Bio, San Jose, CA) and used in RT-PCR with primers LN1036/37. RT-PCRs with five out of six tested seed samples derived from different alfalfa cultivars (Maverick, SW-9215, SW-8421, SW-9720, and CUF101) were virus-positive, indicating a high rate of seed infection (Fig. 2). Resultant amplicons were sequenced and confirmed to be SRAV. Seeds of one cultivar, Regency SY, were RT-PCR-negative (not shown). To additionally confirm seed transmission of the virus, leaves of the germinated seedlings were randomly checked by RT-PCR one week after germination. Except for Regency SY, seedlings of other tested cultivars were positive for SRAV (not shown). These experiments demonstrated localization of SRAV in the internal parts of the seed, likely in the embryo. They also showed a high rate of seed infection by the virus, and its efficient vertical transmission via seeds, thus confirming persistent nature of the virus (Roossinck, 2010).

Dahan et al. (2022) detected SRAV in western flower thrips and suggested a possible role for the insect in virus transmission. Thrips are known to transmit tospoviruses and plant viruses in the *Ilarvirus, Carmovirus, Sobemovirus* and *Machlomovirus* genera (Jones, 2005), whereas endornaviruses are transmitted vertically, through seed via both ova and pollen (Fukuhara and Gibbs, 2012). Although alfalfa is one of the primary hosts for western flower thrips (and other species) and accruing the SRAV during feeding cannot be excluded, transmission of the virus by thrips would require additional experimental confirmation.

Other viruses found in samples infected with SRAV-WA1 included alfalfa mosaic virus, pea streak virus, lucerne transient streak virus, bean leaf roll virus, partitiviruses, and amalgavirus. Fungal and bacterial pathogens, described in alfalfa, like *Alternaria* spp., *Bipolaris* spp., *Stemphylium* spp., *Fusarium* spp., *Pseudomonas* spp., *Erwinia* spp. etc. were also detected. These findings suggested that traditional Koch’s postulate of “one microbe - one disease” should be broadened into the principle of a pathobiome, when disease symptoms are attributed to a diverse community of pathogenic organisms within the plant, rather than to a single infectious agent (Vayssier-Taussat et al. 2014).

Overall, our research confirmed association of SRAV with alfalfa and, for the first time, identified an extensive occurrence of this virus in Washington State. The importance of this work also relies on the hypothesis that SRAV is a persistent endornavirus and its placement within the flavi-like lineage, as suggested by Dahan et al. (2022), may not be entirely accurate. The relationship of the SRAV to endornaviruses appears more likely because of its abundance in alfalfa plants, similar genome organization, seed-mediated transmission, and, at least partly, phylogenetic reconstruction.

More data are needed to assess etiology of the virus and its economic importance. Generally, endornaviruses are associated with symptomless infections and no pathogenic effects (Fukuhara, 2019). *Vicia faba endornavirus* is the only plant endornavirus that leads to the visible phenotype, a cytoplasmic male sterility in the “447” strain of broad bean (*Vicia faba* L.) (Fukuhara, 2019). Interestingly, variations in physiological traits, such as faster seed germination, longer radicle, lower chlorophyll content, higher carotene content, longer pods, and higher weight of 100 seeds, have also been reported in endornavirus-infected common bean lines (Khankhum and Valverde 2018), suggesting that endornaviruses may have symbiotic properties and potential beneficial roles in their hosts (Roossinck, 2015).

The complete genomic sequence of the SRAV (SRAV-WA1 isolate) has been deposited in GenBank under the accession number OP321578.

## Acknowledgements

This study was supported by the United States Department of Agriculture, Agricultural Research Service, CRIS numbers 8042-21000-300-00-D (LGN) and 2090-21000-026-000-D (BMI), and partially by the National Plant Disease Recovery System (NPDRS) grant to LGN.

## Literature Cited

1. Dahan, J., Wolf, Y.I, Orellana, G.E., Wenninger, E.J., Koonin, E.V., and Karasev, A.V. 2022. A Novel Flavi-like Virus in Alfalfa (Medicago sativa L.) Crops along the Snake River Valley. Viruses. 14:1320. doi.org/10.3390/v14061320.

2. Fukuhara, T. 2019. Endornaviruses: persistent dsRNA viruses with symbiotic properties in diverse eukaryotes. Virus Genes. 55:165–173. doi: 10.1007/s11262-019-01635-5.

3. Fukuhara, T., and Gibbs, M.J. 2012. Virus Taxonomy: Ninth Report of the International Committee on Taxonomy of Viruses. 519–521. Elsevier Inc. https://doi.org/10.1016/B978-0-12-384684-6.00048-3.

4. Jones, D.R. 2005. Plant Viruses Transmitted by Thrips. European Journal of Plant Pathology 113:119–157. DOI 10.100.

5. Khankhum, S., and Valverde, R.A. 2018. Physiological traits of endornavirus-infected and endornavirus-free common bean (Phaseolus vulgaris) cv Black Turtle Soup. Arch Virol.163:1051–1056. doi: 10.1007/s00705-018-3702-4.7/s10658-005-2334-1.

6. Kumar, S., Stecher, G., and Tamura, K. 2016. MEGA7: Molecular evolutionary genetics analysis version 70 for bigger datasets. Mol. Biol. Evol. 33, 1870–1874.

7. Meleshko,D., Mohimani, H., Tracanna, V., Hajirasouliha, I., Medema, M.H., Korobeynikov, A., and Pevzner, P.A. 2019. BiosyntheticSPAdes: reconstructing biosynthetic gene clusters from assembly graphs. Genome Res. 29:1352–1362. doi: 10.1101/gr.243477.118.

8. Nemchinov, L.G., and Grinstead S. 2020. Identification of a Novel Isolate of Alfalfa virus S from China Suggests a Possible Role of Seed Contamination in the Distribution of the Virus. Plant Dis. 104:3115–3117. doi: 10.1094/PDIS-04-20-0906-SC.

9. Roossinck, M.J. 2010. Lifestyles of plant viruses. Philos Trans R Soc Lond B Biol Sci. 365: 1899–1905. doi: 10.1098/rstb.2010.0057.

10. Roossinck, M.J., Sabanadzovic, S., Okada,R., and Valverde, R.A. 2011. The remarkable evolutionary history of endornaviruses. J. Gen. Virol. 92(Pt 11):2674–2678. doi: 10.1099/vir.0.034702-0.

11. Roossinck, M.J., 2015. A new look at plant viruses and their potential beneficial rolesin crops. Mol. Plant Pathol. 16, 331–333, DOI: 10.1111/mpp.12241.

12. Shen, X.X., Li, Y., Hittinger, C.T., Chen, X.X., and Rokas A. 2020. An investigation of irreproducibility in maximum likelihood phylogenetic inference. Nature Comm. 11: 6096.

13. Valverde, R.A., Khalifa, M.E., Okada, R., Fukuhara, T., Sabanadzovic, S., and ICTV Report Consortium. 2019. ICTV Virus Taxonomy Profile: Endornaviridae. J. Gen Virol. 100:1204–1205. doi: 10.1099/jgv.0.001277.

14. Vayssier-Taussat M, Albina E, Citti C, Cosson JF, Jacques MA, Lebrun MH, Le Loir Y, Ogliastro M, Petit MA, Roumagnac P, and Candresse T. 2014. Shifting the paradigm from pathogens to pathobiome: new concepts in the light of meta-omics. Front. Cell Infect. Microbiol. 4:29. https://doi.org/10.3389/fcimb.2014.00029.

